# Species invasions shift microbial phenology in a two-decade freshwater time series

**DOI:** 10.1101/2022.08.04.502871

**Authors:** Robin R. Rohwer, Riley J. Hale, M. Jake Vander Zanden, Todd R. Miller, Katherine D. McMahon

**Affiliations:** Department of Integrative Biology, The University of Texas at Austin, Austin, TX, USA; Department of Civil and Environmental Engineering, University of Wisconsin-Madison; Madison, WI, USA; Center for Limnology, University of Wisconsin-Madison; Madison, WI, USA; Zilber School of Public Health, University of Wisconsin-Milwaukee; Milwaukee, WI, USA; Department of Bacteriology, University of Wisconsin-Madison; Madison, WI, USA

**Author notes:** Robin R. Rohwer, **Email:**. **Author Contributions:** Conceptualization: RRR, KDM; Funding acquisition: RRR, TRM, KDM; Methodology: RRR, TRM, JVZ; Investigation: RRR, RJH; Data curation: RRR; Formal analysis: RRR; Visualization: RRR; Writing – original draft: RRR; Writing – review & editing: RRR, RJH, JVZ, TRM, KDM. **Competing Interest Statement:** Authors declare that they have no competing interests.

**Keywords:** Invasive Species, Microbial Ecology, Phenology, Limnology, Time Series

## Abstract

Invasive species impart abrupt changes on ecosystems, but their impacts on microbial communities are often overlooked. We paired a 20-year freshwater microbial community time series with zooplankton and phytoplankton counts, rich environmental data, and a 6-year cyanotoxin time series. We observed strong microbial phenological patterns that were disrupted by the invasions of spiny water flea (*Bythotrephes cederströmii*) and zebra mussels (*Dreissena polymorpha*). First, we detected shifts in *Cyanobacteria* phenology. After the spiny water flea invasion, *Cyanobacteria* dominance crept earlier into clearwater; and after the zebra mussel invasion, *Cyanobacteria* abundance crept even earlier into the diatom-dominated spring. During summer, the spiny water flea invasion sparked a cascade of shifting diversity where zooplankton diversity decreased and *Cyanobacteria* diversity increased. Second, we detected shifts in cyanotoxin phenology. After the zebra mussel invasion, microcystin increased in early summer and the duration of toxin production increased by over a month. Third, we observed shifts in heterotrophic bacteria phenology. The *Bacteroidota* phylum and members of the acI *Nanopelagicales* lineage were differentially more abundant. The proportion of the bacterial community that changed also differed by season; the spring and clearwater bacterial communities changed most following the spiny water flea invasion that lessened clearwater duration and intensity, while the diverse summer bacterial community changed least following the zebra mussel invasion despite the observed shifts in diversity and toxicity during summer. These long-term invasion-mediated shifts in microbial phenology demonstrate the interconnectedness of microbes with the broader food web, and their susceptibility to long-term environmental change.

**Significance Statement:** Microbial communities are typically studied as part of the microbial loop, separately from the broader food web. Using a two-decade freshwater time series, we explored whether two species invasions that shifted the metazoan food web (spiny water flea and zebra mussels) also impacted the microbial communities. We looked for seasonal responses because the microbial communities had strong seasonal patterns. We discovered that *Cyanobacteria* increased early in the year, and *Cyanobacteria* diversity increased in the summer. Cyanotoxins also increased, along with the duration of toxin production. In the heterotrophic bacterial community, some organisms changed consistently within lineages and seasons while others diverged. These findings illustrate the importance of seasonal context, and highlight the interconnectedness of bacteria with the broader food web.

## Introduction

Invasive species have wide-ranging impacts on ecosystems, often rewiring food webs and disrupting nutrient cycling, and they can be even more disruptive than global abiotic change (1). However, the impact of species invasions on entire microbial communities are rarely considered. Microbes are usually considered separately from metazoan lake ecology; bacterial communities are thought to interact with the broader food web as part of a microbial loop, where phytoplankton-derived dissolved organic matter is returned to higher trophic levels (2, 3). Bacteria that consume organic matter are in turn consumed by ciliates and nanoflagellates, which are eaten by microzooplankton, thus linking microbial communities with higher trophic levels. Although this microbial food web is often studied separately from the metazoan food web, we hypothesized that metazoan species invasions would change the bacterial community.

We collected a 20-year microbial community time series (2000-2019, 496-samples) from Lake Mendota, a eutrophic temperate lake in WI, USA. This period includes two metazoan species invasions: the predatory zooplankton spiny water flea in 2009 (*Bythotrephes cederströmii*) and zebra mussels in 2015 (*Dreissena polymorpha*). The spiny water flea invasion triggered a food web cascade that resulted in a one meter loss in water clarity (4), as increased zooplankton predation by spiny water flea led to reduced grazing pressure on phytoplankton. This impact was particularly strong during the spring clearwater phase, which decreased in intensity and duration (5). The invasion of zebra mussels triggered a 300% increase in benthic zooplankton and phytoplankton abundance, and they themselves became abundant enough to filter the epilimnion volume every 45 days (6). Zebra mussels increase filtration, but they can also worsen *Cyanobacteria* blooms via selective retention of eukaryotic algae and release of Cyanobacteria back into the lake (7), and by altering nutrient stoichiometry (8). In Lake Mendota, zebra mussels did not impact the pelagic water clarity in either direction (6). These mid-time series disturbances set up a natural experiment with which we explored the impact of metazoan species invasions on the bacterial community.

Lake Mendota is a temperate, dimictic lake and its ecology follows consistent seasonal patterns (phenology). The lake freezes in the winter, and ice-off is followed by spring mixing and a diatom bloom. As the water warms, zooplankton grazing increases and results in a short-lived period of high water clarity, which ends when planktivorous fish begin eating zooplankton (5). After this clearwater phase, diatoms give way to *Cyanobacteria* which persist throughout the summer (9). During the summer the water column is thermally stratified, and anoxia develops in the bottom layer while nutrient draw-down occurs in the upper layer. When temperatures drop in the fall, Lake Mendota mixes and remains mixed until ice-on. Seasonal patterns have also been observed in microbial community composition (10), and these seasonal cycles can mask long-term changes (11). Therefore, we split our time series into seasons and looked for changes in bacterial phenology.

Here we identify seasons that described the bacterial phenology, and then identify phenological change in *Cyanobacteria*, cyanotoxins, and heterotrophic bacteria. These observed phenological shifts are a direct observation of multi-decadal microbial change in response to invasive species.

## Results

We characterized the microbial community composition via 16S rRNA gene amplicon sequencing performed on duplicate water grabs (961 total samples, 496 sample dates) (Fig 1). The water grabs integrated the top 12 m of the water column, which approximates the mixed upper layer. In addition to the sequencing-based relative abundances, we obtained environmental data available from the North Temperate Lakes Long Term Ecological Research program (NTL-LTER), including water physical and chemical measurements and microscopy-based counts of zooplankton and phytoplankton.

**Figure 1.**
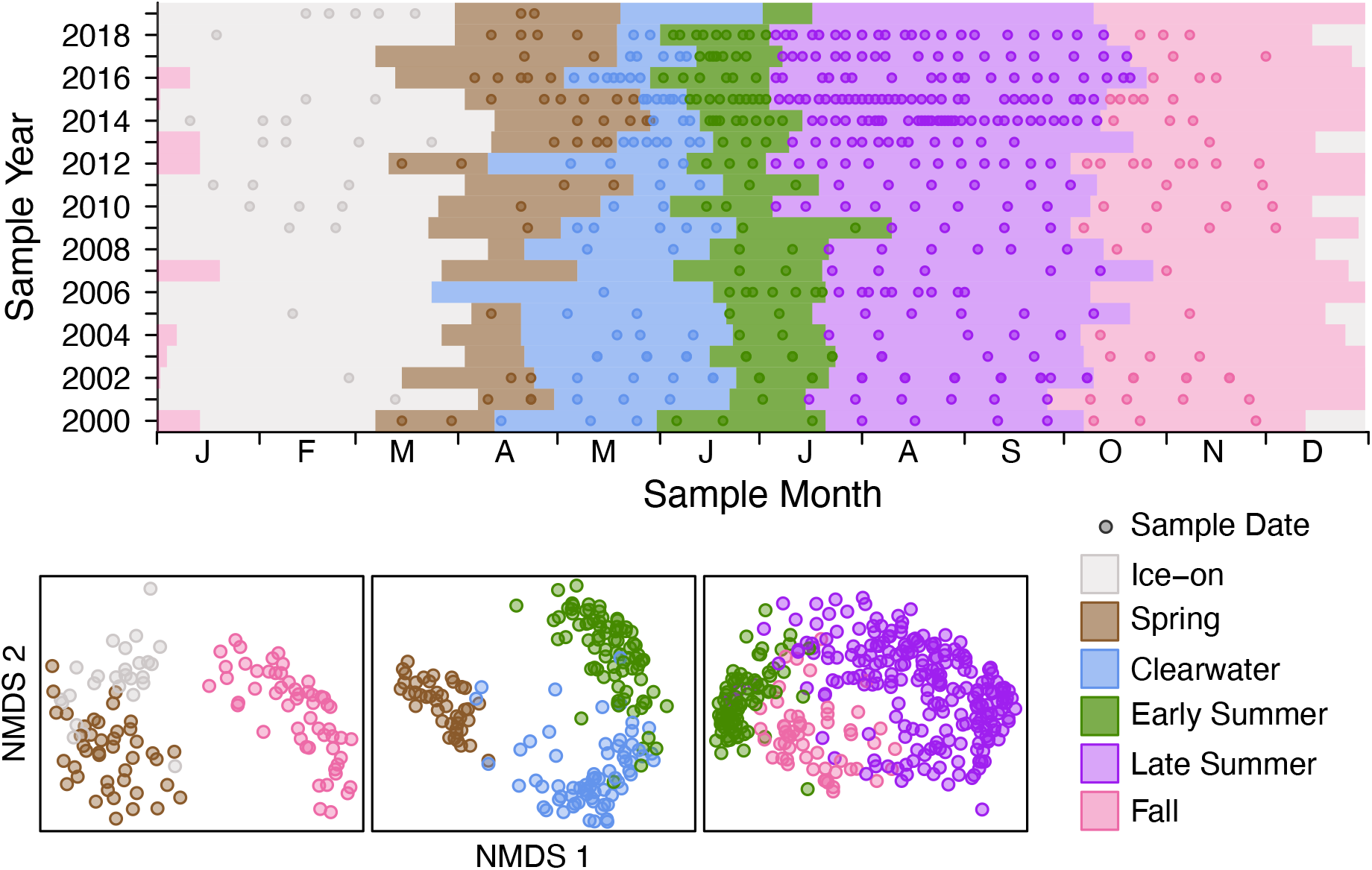
Sample Dates and microbial phenology. **(A)** Sample dates are represented by points, and season durations are represented by shading. **(B)** Non-metric multidimensional scaling of CLR-transformed bacterial community compositions. Points represent sample dates, and colors indicate seasons. Three seasons are compared in each NMDS plot to highlight season boundaries. All NMDS stress values are **≤** 0.1.

### Microbial Phenology

To compare across years, we defined lake seasons using environmental variables and compared samples within the same season. We compared across multiple years by season instead of by Julian date to discriminate shifts in each season’s community composition from shifts in seasonal timing. We created multiple potential season definitions based on environmental data, and then selected the definitions that were most representative of the microbial phenology. We applied non-metric multidimensional scaling (NMDS) (12) to sample distance matrices calculated with the phi distance metric for centered log-ratio (CLR)-transformed relative abundances (13). To select the season definitions most relevant to the microbial community, we chose definitions that resulted in clear separation between season boundaries in NMDS plots (Fig 1). Samples grouped by our selected season definitions were significantly less variable within groups than between groups for every season boundary (ANOSIM < 0.001).

The season definitions that best described the microbial phenology were ice-on, spring, clearwater, early summer, late summer, and fall (Fig 1, Table S1). The ice-on season was determined by the onset and break-up of continuous ice-cover across a central lake transect. Spring began with ice-off and ended with clearwater onset, spanning a period that includes the spring diatom bloom (9). Clearwater was defined based on water clarity measurements, and is an annual phase driven by high zooplankton grazing pressure (5). Early summer was a period of growing stratification, bounded by clearwater and the point at which hypolimnetic anoxia extended above 12 m. Late summer was a period of strong stratification and anoxia, which ended with fall mixing. We defined mixing as a difference in average epilimnion and hypolimnion temperatures < 3°C. Fall began with mixing and ended with ice-on.

### Shifts in *Cyanobacteria* phenology

Given spiny water flea’s marked disruption of clearwater phenology, *i*.*e*. shortened duration and intensity, we first looked for changes in spring phytoplankton phenology. Historically, the end of clearwater phase marks the transition from a diatom-dominated system to a *Cyanobacteria*-dominated system (9). We found that after the spiny water flea invasion, *Cyanobacteria* began appearing during clearwater phase (Fig 2A). We did not see a similar increase in chloroplast relative abundance or diatom counts during clearwater phase, indicating that the transition to *Cyanobacteria* dominance shifted earlier. We further found that after the zebra mussel invasion, *Cyanobacteria* began appearing even before the start of clearwater phase (Fig 2A). Count data confirmed that the biovolume of *Cyanobacteria* increased 2-fold, growing to over 10% of total phytoplankton (Fig S1). This earlier onset of *Cyanobacteria* describes a kingdom-level regime shift from a eukaryotic algae-dominated period to one that includes *Cyanobacteria*.

**Figure 2.**
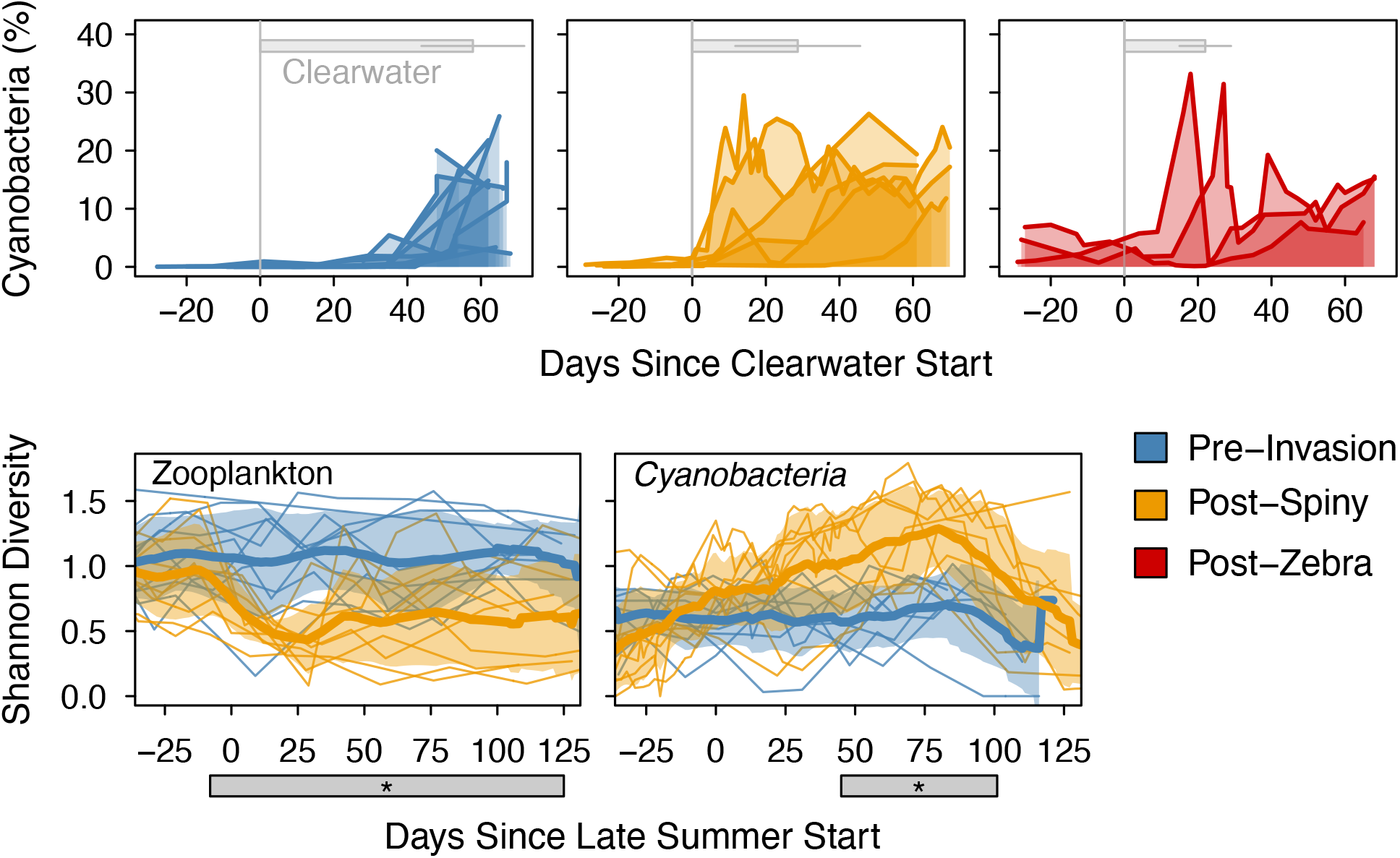
Shifts in *Cyanobacteria* phenology. **(A)** Spring relative abundance of *Cyanobacteria* aligned as days since clearwater. Average and standard deviation duration of clearwater is indicated by grey bars. Blue represents the pre-invasion period of 2000-2009, orange the post-spiny water flea period of 2010-2015, and red the post-zebra mussel period of 2016-2018. **(B)** Summer diversity of zooplankton and *Cyanobacteria*. Thick lines represent the mean Shannon diversity and shading represents the standard deviation. Thin lines represent individual years. Here the orange post-spiny water flea period includes years 2010-2018. Grey starred regions indicate the period of significant difference in mean diversity (p < 0.05). *Cyanobacteria* diversity is shown at the order level, and Zooplankton diversity is shown at ecologically relevant groupings with spiny water flea counts removed.

After identifying these changes in the spring phenology, we next looked for spiny water flea invasion impacts on the summer cyanobacterial phenology. *Cyanobacteria* abundance (relative to the microbial community) during summer and fall did not change with the spiny water flea invasion. However, *Cyanobacteria* diversity during the late summer season increased (Fig 2B). This increase in cyanobacterial diversity was observed at fine and coarse-resolution taxonomic levels, including levels as coarse as order (Fig 2B). It is possible that the shift in summer diversity could be explained by the earlier onset of the *Cyanobacteria-*dominated season resulting in an earlier onset of high *Cyanobacteria* diversity. However, the spring cyanobacterial incursion was comprised of a single amplicon sequence assigned to *Aphanizomenon*, not to a more complex community typical of early summer. Additionally, we observed a concurrent late summer change in zooplankton diversity; after the spiny water flea invasion, zooplankton diversity decreased (Fig 2B, Table S2). Therefore, we attribute the shift in cyanobacterial summer diversity to a cascade of shifting diversity originating in the zooplankton community. The lower diversity of zooplankton may have resulted in narrower grazing pressure, allowing a wider range of *Cyanobacteria* to flourish.

Next, we looked for changes in the summer cyanobacterial community stemming from the zebra mussel invasion. We first compared total *Cyanobacteria* abundance before and after zebra mussels but did not see strong and consistent changes in relative abundance or biovolume (Fig S1). We also did not observe strong shifts in cyanobacterial community composition. *Cyanobacteria* can produce potent toxins, but *Cyanobacteria* abundance does not always correlate with cyanotoxin concentrations (14, 15), suggesting that toxin concentrations could yet have changed as a result of the invasion.

### Shifts in cyanotoxin phenology

To examine the impact of zebra mussels on cyanobacterial toxicity, we collected a 6-year, 246-sample toxin time series spanning the 3 years preceding and following the zebra mussel invasion. We measured the hepatotoxin microcystin and the bioactive peptide anabaenopeptin using high performance liquid chromatography coupled with tandem mass spectrometry (HPLC-MS/MS) (Table S3) (16). We observed a 3-fold decrease in anabaenopeptin and a concurrent 3-fold increase in microcystin during early summer (p < 0.05) (Fig 3A). The family *Microcystaceae* is considered the main microcystin producer in Lake Mendota, although it is also capable of producing anabaenopeptin (14). However, the relative abundance of *Microcystaceae* did not change. This suggests that the switch from anabaenopeptin to microcystin is due to more complex ecological drivers than the abundance of *Microcystaceae*, and illustrates the complexity of invasion impacts. This shift in toxin concentrations has implications for human health, because microcystin is a more potent toxin than anabaenopeptin (17).

**Figure 3.**
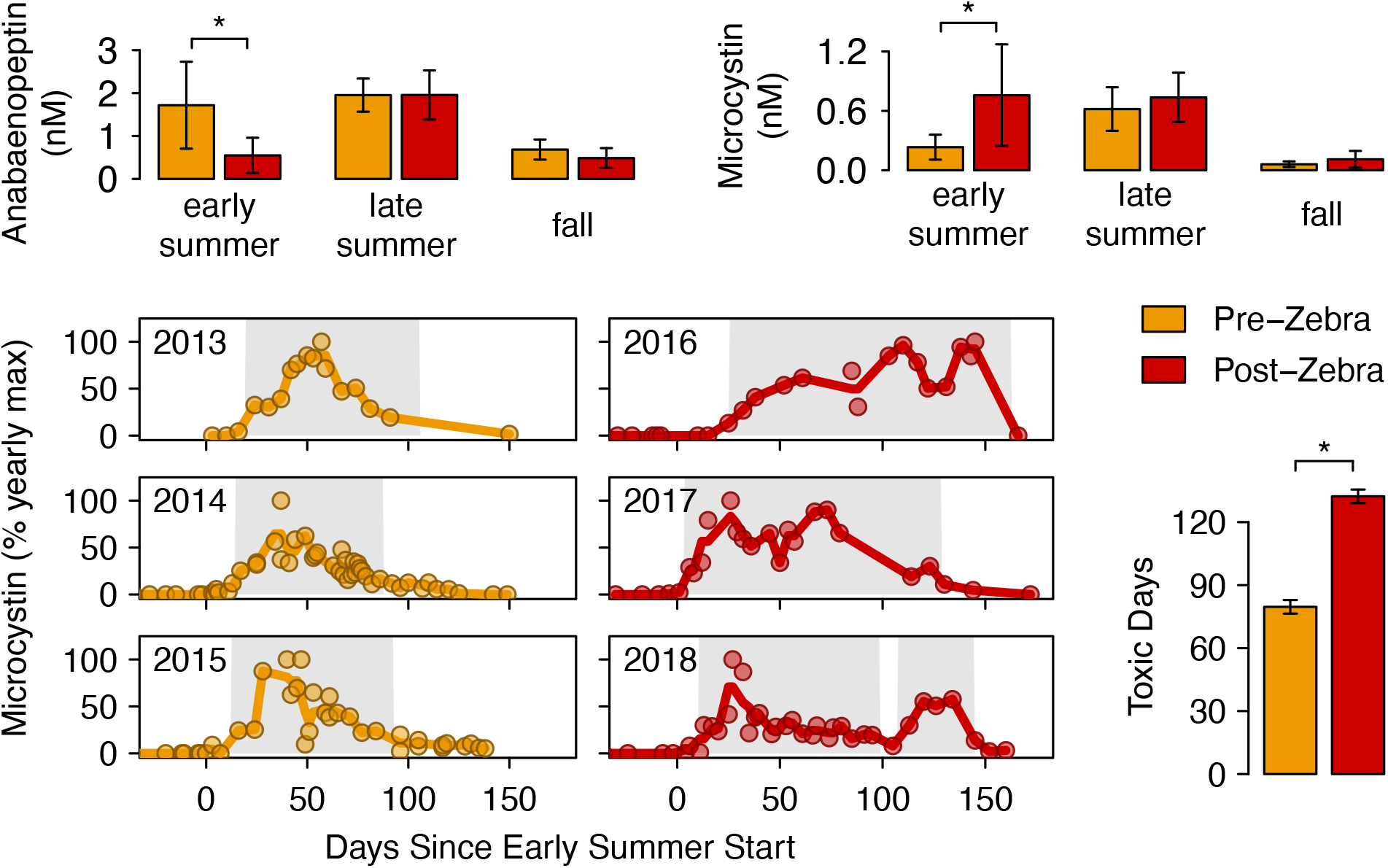
Shifts in cyanotoxin phenology. Orange represents the pre-zebra mussel invasion period of 2013-2015, red represents the post-zebra mussel invasion period of 2016-2018. Stars represent significant differences (p < 0.05). **(A)** Concentration of ananbaenopeptin and microcystin during the seasons with regular toxicity. **(B)** Microcystin toxin concentrations normalized by the yearly maximum concentrations to compare annual patterns. Shaded regions represent the duration of toxin production, defined as days when toxins were present at ≥ 15% of maximum toxicity. **(C)** Toxin production duration compared before and after the invasion of zebra mussels.

After observing this increase in early summer microcystin, we next explored the phenology of microcystin production (Fig 3B). Total toxin concentrations were variable between years, so for each year we normalized total microcystin by the maximum measured concentration within that year. We found that the duration of toxin production, defined as the period during which toxins were at least 15% of maximum annual toxicity, increased after the zebra mussel invasion by 53 ± 9 days, or 170 ± 10%. (p < 0.001) (Fig 3C). This increase was due to a longer late summer season, as well as toxin production extending further into the early summer and fall seasons. We observed these increases in toxicity despite the lack of change in the *Cyanobacteria* abundance and diversity. This might be explained by changes in gene expression, which can be variable enough that the abundance of microcystin *mcy* genes is a poor predictor of microcystin concentrations (14). Another possible explanation is preferential retention of non-toxic strains or individual cells, with toxic cells released back into the water column. Zebra mussels were previously found to preferentially retain a non-toxic *Microcystis* strain over a toxic one (18).

In summary, cyanobacterial phenology shifted in response to the two metazoan invasions, with changes in timing, diversity, and toxicity.

### Shifts in heterotrophic bacteria phenology

To explore whether invasion impacts extended further, into the heterotrophic bacterial community, we compared each season’s heterotrophic community before and after invasion using NMDS analyses of their CLR-transformed phi dissimilarities (13). We observed modest grouping based on invasion status (Fig S2), and an ANOSIM analysis confirmed that within-group differences were smaller than between-group differences in all seasons (ANOSIM significance = 0.001).

To identify which bacteria were responsible for the shift in community composition, we performed a differential abundance analysis after applying a CLR transformation to account for the relativized nature of sequencing data (19). We used a generalized linear model to test for organisms that were differentially abundant before and after each invasion (20), and we looked for changes within each season. We considered an operational taxonomic unit (OTU) differentially abundant when it had an effect size > 0.5, and we manually confirmed the ecological significance of this effect size cutoff by inspecting relative abundance over time (examples in Fig 4).

**Figure 4.**
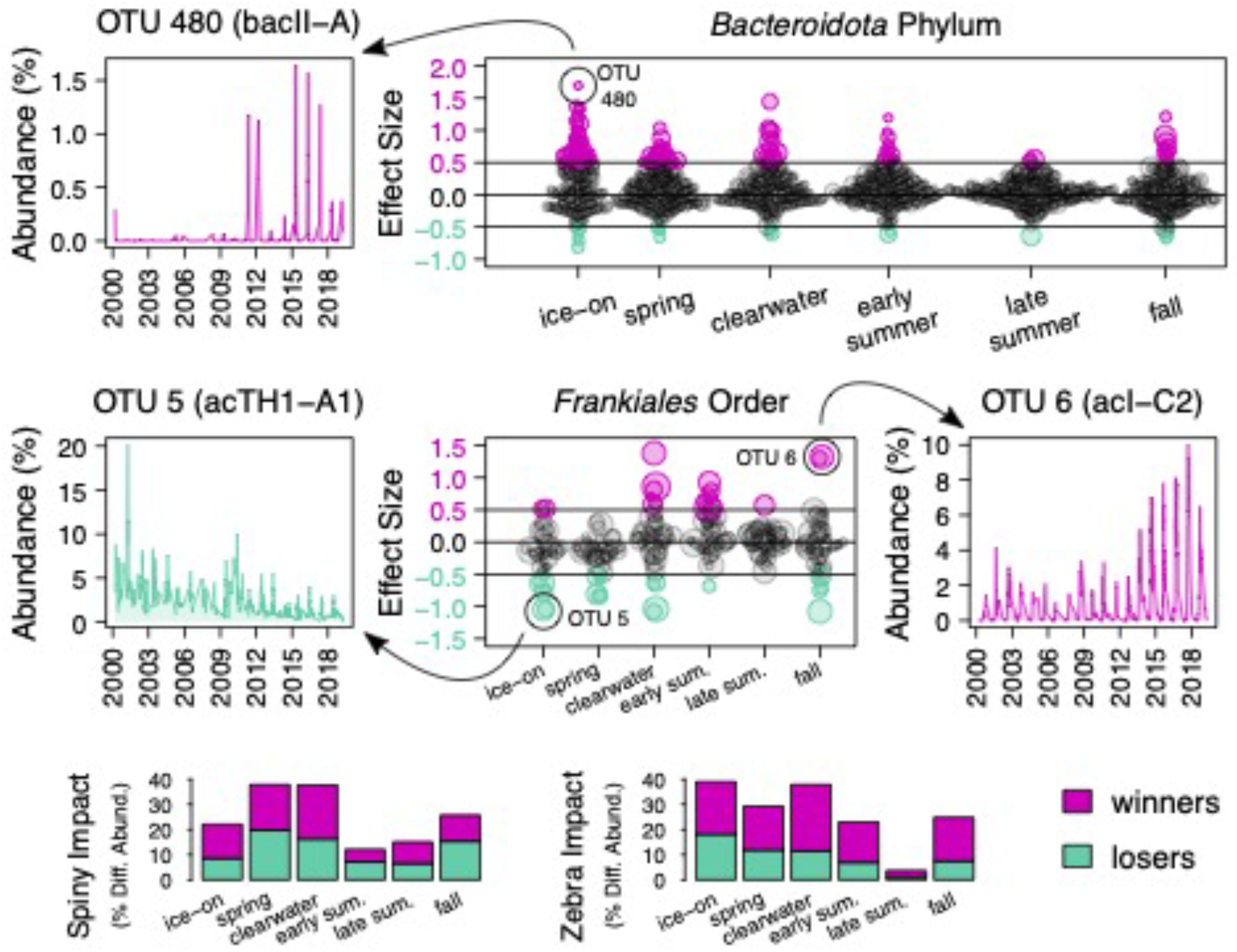
Shifts in heterotrophic bacteria phenology. Differential abundance was calculated on CLR-transformed abundances with a GLM model. Pink represents “winners” that differentially increased, and teal represents “losers” that differentially decreased. Points represent OTUs, and point sizes are log-scaled by the OTU’s mean abundance. **(A)** Left, an example relative abundance over time curve of an OTU with positive effect size. Right, effect sizes for differential abundance of phylum *Bacteroidota* OTUs around the zebra mussel invasion. **(B)** Middle, effect sizes for differential abundance of order *Frankiales* OTUs around the zebra mussel invasion. Right and left are example winner and loser OTU abundances over time. **(C)** Percent of each season’s microbial community that was differentially abundant over time. OTUs were normalized by their mean abundance before calculating the percentage of community change. The impact of the spiny water flea invasion is shown on the left, and the impact of the zebra mussel invasion is shown on the right.

To assess which seasonal communities were most susceptible to change, we looked at the proportion of OTUs within the community that were differentially abundant. To better reflect the ecological impact of the differentially abundant OTUs, we normalized the OTUs by their mean relative abundance. The invasions impacted seasonal community composition to different degrees. For example, spiny water flea had the largest impact on the proportion of the community that changed during spring and clearwater, with 38% of the average community becoming differentially abundant (Fig 4C, left). This means that in addition to changing the seasonal timing of clearwater (5), spiny water flea also changed the heterotrophic bacteria community. Meanwhile, zebra mussels had the smallest impact on the late summer microbial community, with only 4% of the average community becoming differentially abundant (Fig 4C, right). The late summer microbial community had the smallest proportion of differentially abundant organisms, and also the fewest overall differentially abundant OTUs (Fig 4). The late summer community also has the highest alpha diversity, and high biodiversity has been linked to ecosystem stability (21). It is possible that this high diversity makes the late summer ecosystem more robust. The seasonal differences in invasion impacts emphasize the importance of a phenological context.

Among the organisms that were differentially abundant, OTUs in the phylum *Bacteroidota* were zebra mussel invasion “winners” broadly (Fig 4A, Table S4). These winners included abundant OTUs from the bacI, bacII, bacIII, and bacIV lineages (*Chitinophagales, Flavobacteriales*, and *Cytophagales* orders, lineage nomenclature from (22, 23)). However, most changes in differential abundance were not consistent within taxonomic lineages or across seasons. For example, within the abundant lineages of order *Frankiales, Nanopelagicaceae* (acI) tended to win while acTH1 and acSTL tended to lose (Fig 4B, Table S4).

The *Bacteroidota* phylum is often particle-associated (23, 24), and *Bacteroidota* OTUs were broadly differentially abundant after both the spiny water flea invasion and the zebra mussel invasion. In contrast, the *Frankiales* order is comprised of small, free-living bacteria and contained both winner and loser OTUs. The non-uniform response of the *Frankiales* order suggests that they occupy divergent niche spaces. Further, it appears that the niche spaces for acI may be seasonally dependent. The acI lineage are oligotrophs (25), and acI-A and acI-B tended to win during clearwater and early summer even though they are abundant throughout the year, while acI-C is abundant during fall and also tended to win during fall. AcTH1 and acSTL are also small, free-living *Frankiales*, but little is known about their ecology and metabolism. They comprised many of the abundant invasion losers in all seasons after both invasions, which suggests that their niche is less seasonally dependent and different from their close relatives’.

In summary, heterotrophic bacteria phenology was shifted by metazoan species invasions. Some organisms showed consistent responses within phyla and across seasons, and some organisms showed divergent responses within orders and between seasons, suggesting that some niche spaces were changing broadly while others experienced seasonally dependent changes. Overall, the amount of change bacterial communities experienced was seasonal, and some seasonally-defined communities were more robust to change than others.

### Species invasions are drivers of change

To compare the importance of species invasions to other environmental drivers, we evaluated metrics related to invasion status, landscape change, and climate change in a modeling framework. We predicted the lag between *Cyanobacteria* onset and clearwater (Fig 2A) and the CLR-transformed abundance of the winner acI-B OTU 1 (Fig 4B) using invasion status, phosphorus loading, ice duration, and hot summer days. We used Akaike Information Criterion (AIC) to identify which drivers were included in the best-performing models (Table S5 and S6). Spiny water flea and zebra mussel invasion status were included in all the best models, which emphasizes the importance of species invasions as drivers of changing microbial phenology. The number of hot summer days and mean phosphorus loading was also included in some of the best-performing models. This suggests that even though species invasions were a strong driver of microbial change during the 19-year time frame of our dataset, climate and landscape are interacting drivers of microbial change.

## Discussion

Over century-long time frames, the interacting driver of climate change is shifting the phenology of northern hemisphere lakes. Winters are shortening (26), summer stratification and anoxia is increasing (27), and extreme weather is becoming more common (28). In Lake Mendota specifically, ice duration (29) and clearwater (5) are shortening, while the stratified (29) and anoxic (30) periods are lengthening. We posit that lake microbial communities will also change in response to climate change because the lake microbial community has phenological patterns that track these shifting seasons. Further, we directly observed that food web-mediated shifts in seasonal timing can change microbial community timing, composition, and toxicity. It follows that the interacting driver of climate change shifting seasons over longer time frames will likely shift microbial phenology too.

Moreover, the spiny water flea invasion itself was driven by climate change. Previous work on Lake Mendota found that an unusually cool year allowed a sleeper population of spiny water flea to irrupt (31), and weather extremes like this are expected to increase with climate change (28). It is possible that the spiny water flea-mediated food web shift also paved the way for the zebra mussel invasion. Zebra mussels likely had multiple introductions to Lake Mendota based on the large amount of recreational boat traffic on the lake, so it is plausible that they were introduced prior to 2015 and existed as a low abundance sleeper population. The establishment of one invasive species can destabilize an ecosystem in a way that primes it for additional invasions (32), or that triggers the population irruption of existing sleeper populations (33). A climate anomaly triggered a population irruption of the invasive spiny water flea sleeper population, and the resulting food web disruptions may have paved the way for the establishment of zebra mussels. The impact of these invasions reached all the way into the bacterial community, illustrating how the interacting drivers of species invasions and climate change can have complex and far-reaching impacts.

Harmful cyanobacterial blooms are predicted to increase with warmer temperatures, higher CO_2_, and increasingly intense rain events (34). We observed an earlier onset of Cyanobacteria into the clearwater and spring seasons. In other lakes, regime changes have been observed as oligotrophic systems switch to a new stable state dominated by *Cyanobacteria* (35). It is possible that we may be observing the beginning of a transition to a new stable state at a seasonal level. In the summer seasons already dominated by *Cyanobacteria*, we found the fewest differentially abundant organisms. This suggests that even if tipping points are reached in other seasons, the community is less likely to shift when it would be most desired: during the summer months plagued by toxic cyanobacterial blooms.

Long-term change is difficult to distinguish from short-term variability and seasonal cycles, a phenomenon termed the “invisible present” (11). Observations of the microbial community for almost twenty years have allowed us to distinguish consistent phenological patterns, and to identify separate long-term, invasion-mediated shifts. The establishment and continuation of time series like ours are the only way to directly observe the long-term impacts of environmental change. Our study found that microbial communities are sensitive to long-term change, specifically to the cascading impacts of invasive species. These impacts included many aspects of phenology, from timing and abundance, to diversity and community composition, to human health and cyanobacterial toxicity.

## Materials and Methods

### Bacterial Community Data

Microbial sampling, sequencing, and data processing was performed as described in detail in Rohwer & McMahon (36). Briefly, integrated epilimnion samples were collected from the central deep hole of Lake Mendota (WI, USA) using a 12 m tube, and water was well mixed before filtering onto a 0.22 um filter (Pall Corporation). Filters were stored in a -80°C freezer until DNA extraction in 2018-2019. DNA was extracted using a FastDNA Spin Kit (MP Biomedicals) after sample order was randomized. Samples were sent to Argonne National lab for amplification of the 16S rRNA gene V4-V5 region (515F-Y GTGYCAGCMGCCGCGGTAA, 926R CCGYCAATTYMTTTRAGTTT) and sequencing on an Illumina MiSeq instrument.

After sequencing, data was processed using custom R scripts available at https://github.com/rrohwer/limony and described in Rohwer & McMahon (36). Microbial relative abundances from replicate samples were averaged, and a standard deviation was calculated. This processed data is available as part of an R package at https://github.com/rrohwer/limony. The raw sequencing data is available through the NCBI Sequence Read Archive under accession number PRJNA846788.

### Environmental Data

We collected lake physical and chemical data alongside microbial samples, including water temperature and dissolved oxygen measured with a YSI ProPlus multiparameter sonde (Yellow Springs Instruments), and water clarity measured with a Secchi disk. This data is available through the Environmental Data Initiative (EDI) portal under identifiers knb-lter-ntl.415.2 (37) and knb-lter-ntl.416.1 (38). We combined our measurements with additional water temperature, dissolved oxygen, water clarity, and ice cover datasets made available by the North Temperate Lakes Long-Term Ecological Research program (NTL-LTER) under EDI identifiers knb-lter-ntl.29.29 (39), knb-lter-ntl.130.29 (40), knb-lter-ntl.335.1 (41), knb-lter-ntl.400.2 (42), knb-lter-ntl.31.30 (43), and knb-lter-ntl.33.35 (44). We obtained mean annual phosphorus loading at the Yahara River Windsor site from the USGS surface water annual statistics (45).

We defined lake seasons using metrics calculated from this environmental data. We identified the epilimnion, metalimnion, and hypolimnion depths by identifying the inflection points of temperature profile vectors, calculated the average epilimnion and hypolimnion temperatures for all complete profiles, and smoothed these values with a 7-day moving average. We defined mixing as the day when the linearly interpolated difference in epilimnion and hypolimnion temperature fell below 3°C. We considered depths with < 1 mg/L dissolved oxygen anoxic, and we applied a 7-day moving average to the depth of the anoxic layer. We defined the start of the late summer season as the day when the linearly interpolated anoxic layer depth reached 12 m. We defined the clearwater season’s start and end for each year by manually identifying the edges of the abrupt drop in Secchi depth measurements that occurs after ice-off, and we refined the edge dates based on chloroplast and *Cyanobacteria* abundance when sequencing data existed without a paired Secchi depth measurement.

We obtained absolute abundance and biovolume of large *Cyanobacteria* and eukaryotic algae using microscopy-based phytoplankton counts from the NTL-LTER, and we used microscopy-based zooplankton counts to calculate zooplankton diversity. This data is available under EDI identifiers knb-lter-ntl.88.30 (46) and knb.lter.ntl.90.31 (47).

### Cyanotoxin Data

We measured toxins in 246 whole-water samples spanning 6 years (2013-2018). These samples were taken from the same water grabs used to characterize the microbial community. We quantified cyanotoxins using methods described in detail by Miller *et al*. (16). Briefly, lyophilized samples were resuspended in formic acid, subjected to three freeze-thaw cycles, and extracted in methanol and formic acid in a sonicating water bath. The extract supernatant was analyzed by targeted high performance liquid chromatography coupled with tandem mass spectrometry (HPLC-MS/MS).

Eleven microcystin (MC) congeners and three anabaenopeptin (Apt) congeners were analyzed. Certified reference standards of MC-LR, [Dha^7^]MC-LR and nodularin were purchased from the National Research Council of Canada Biotoxins program (Halifax, Nova Scotia). MC-LA, MC-RR, MC-LF, MC-YR, MC-WR, MC-LY, MC-LW, MC-HtyR, MC-HilR (all >95%) were purchased from Enzo Life Sciences (Farmingdale, NY). Anabaenopeptin A (Apt-A) (> 95%), Apt-B (>95%), and Apt-F (> 95%) were purchased from MARBIONC (Wilmington, NC). To calculate total concentrations, we summed nanomolar measurements of the congeners. 99 ± 4% of total microcystin was comprised of the congeners MC-LA and MC-LR. Toxin totals are available in a tabular format as Table S2.

### Analysis

We defined microbially-relevant seasons (Fig 1A) based on the environmental data. Ice extent, mixing, and clearwater were phenological markers that corresponded well with the microbial community. To divide the summer months we explored multiple seasonal cutoffs based on stratification extent, epilimnion temperature, and anoxia extent. We visualized each season option with non-metric multidimensional scaling (NMDS) analysis and manually chose the most relevant definitions based on visual inspection of the NMDS plots. To account for the proportional nature of sequencing data, we performed a centered log ratio (CLR) transformation on our data before the NMDS analysis (Fig 1B, Fig S2). Following the guidance provided by Quinn *et al*. (19), we filtered each 3-season group to contain only OTUs that occurred in at least 10% of the samples, assigned small-values to zeros using the zCompositions R package (48), and performed a CLR-transform and calculated the *phi* distance metric using the propr R package (13). We performed NMDS and confirmed the significance of our season choices by calculating the ANOSIM significance using the vegan R package (12). The season start dates are listed in Table S1. We used season start dates to align samples in our subsequent analyses; instead of aligning samples in different years by Julian date, we aligned samples by the days since the start of a given season. Comparisons across Julian dates do not account for shifts in seasonal timing across years, but aligning samples by season start dates enables comparison of phenologically similar microbial communities.

We observed changes in spring *Cyanobacteria* relative abundance (Fig 2A) by aligning samples by the start of clearwater phase and grouping years into invasion periods. Both invasions were discovered in fall, so we considered the subsequent years the start of the invasion period. The seven 2019 samples were excluded from this analysis because they did not span the entire spring-clearwater-early summer period. We compared the summer diversity before and after the spiny water flea invasion (Fig 2B) by aligning samples by the start of the early summer season and calculating the Shannon diversity index using the vegan R package (12). We observed no significant difference in diversity between the spiny water flea-only period and the zebra mussel period, so we grouped these two periods together as a single post-spiny water flea group. Before calculating *Cyanobacteria* diversity, we excluded samples from 2000-2003 that had been prefiltered with a 10 µm filter instead of filtered directly onto a 0.22 µm filter. Before calculating zooplankton community diversity (Fig 2B), we excluded spiny water flea and juvenile copepods from the zooplankton count data and grouped zooplankton based on ecological relevance and our confidence in the identifications (Table S3). We calculated the mean and standard deviation of linearly interpolated daily diversity values and performed a t-test to compare between invasion groups.

We calculated the concentration of toxins before and after the zebra mussel invasion (Fig 3A) by separating toxin measurement dates by season and splitting the years into invasion groups. For each season and year group, we calculated the mean and standard deviation of all toxin measurements within it. We performed a t-test on each seasonal block to identify significant change between invasion groups. We calculated the duration of toxin production (Fig 3B-C) by normalizing microcystin concentrations in each year by the yearly maximum concentration. We applied a linear interpolation to the 7-day moving average of measurements (lines in Fig 3B). We defined days as toxin-producing if their interpolated daily toxin concentration was > 15 % of the yearly maximum value.

We identified differentially abundant bacteria (Fig 4) using the ALDEx2 package (49) following the guidance provided by Quinn *et al*. (19). For each season group, we removed OTUs present in fewer than 10% of samples and added small numbers to remaining zeros using zCompositions (48). We identified differentially abundant organisms and their effect sizes using a generalized linear model with the aldex.glm and aldex.glm.effect functions from the ALDEx2 package (49). We manually chose an effect size cutoff of 0.5 by examining plots of OTU abundance over time for OTUs with varying effect sizes (examples in Fig 4A and Fig 4B). We identified which seasons were most susceptible to microbial community change (Fig 4C) by calculating the proportion of the community that was differentially abundant. We defined this proportion as the average abundance of all differentially abundant OTUs divided by the average abundance of all OTUs. All differentially abundant bacteria are listed in Table S4.

We created generalized linear models to predict microbial phenology from invasion status, Yahara River phosphorus loading into the lake, ice duration, and hot summer days (Table S4-5). We used the glm function in R to create models, and the dredge function in the MuMIn R package to compute the corrected AIC for each combination of predictors. For a *Cyanobacteria* phenology response variable, we modeled the lag between clearwater season start and the first day with *Cyanobacteria* abundance ≥ 3%. For a heterotroph phenology response variable, we modeled the average CLR-transformed clearwater abundance of OTU 1, a *Frankiales* acI-B that was a spiny water flear and zebra mussel clearwater winner and is also the most abundant OTU in the dataset. As predictor variables, we used the previous-winter ice duration as a climate metric (44), the mean annual Yahara River phosphorus loading as a landscape metric (45), and the number of days with an average epilimnion temperature ≥ 23°C as a climate metric related to the initial spiny water flea invasion (31).

## Acknowledgments

We thank Stephen Carpenter and Edward DeLong for friendly reviews. We thank technicians Ame Xiong and Leah Stromberg for assistance measuring cyanotoxins. We thank Jake Walsh for assistance interpreting the zooplankton data. We thank the many field personnel involved in microbial sampling, including sampling leads Angela Kent, Tony Yannarell, Ashley Shade, Stuart Jones, Ryan Newton, Georgia Wolfe, Emily Kara Read, Lucas Beversdorf, and James Mutschler; and initial Microbial Observatory lead Eric W. Triplett.

We thank the funding sources that made this work possible:

U.S. National Science Foundation Postdoctoral Research Fellowship in Biology, award number 2011002 (RRR)

E. Michael and Winona Foster WARF Wisconsin Idea Fellowship, 2018 (RRR)

National Institute of Food and Agriculture, U.S. Department of Agriculture, award number 2016-67012-24709 (Joshua J. Hamilton)

National Institute of Food and Agriculture, U.S. Department of Agriculture, Hatch Projects WIS01516, WIS01789, WIS03004 (KDM)

U.S. National Science Foundation North Temperate Lakes Long-Term Ecological Research site, NTL-LTER, award numbers DEB-9632853, DEB-0217533 (Stephen Carpenter) and DEB-0822700, DEB-1440297 (Emily Stanley)

U.S. National Science Foundation Microbial Observatories program, award numbers MCB-9977903 (Eric Triplett) and DEB-0702395 (KDM)

U.S. National Science Foundation INSPIRE award, DEB-1344254 (KDM)

The Laboratory for Aquatic Environmental Microbiology & Chemistry at the University of Wisconsin-Milwaukee, Zilber School of Public Health (TRM)

---

**Figure S1.**
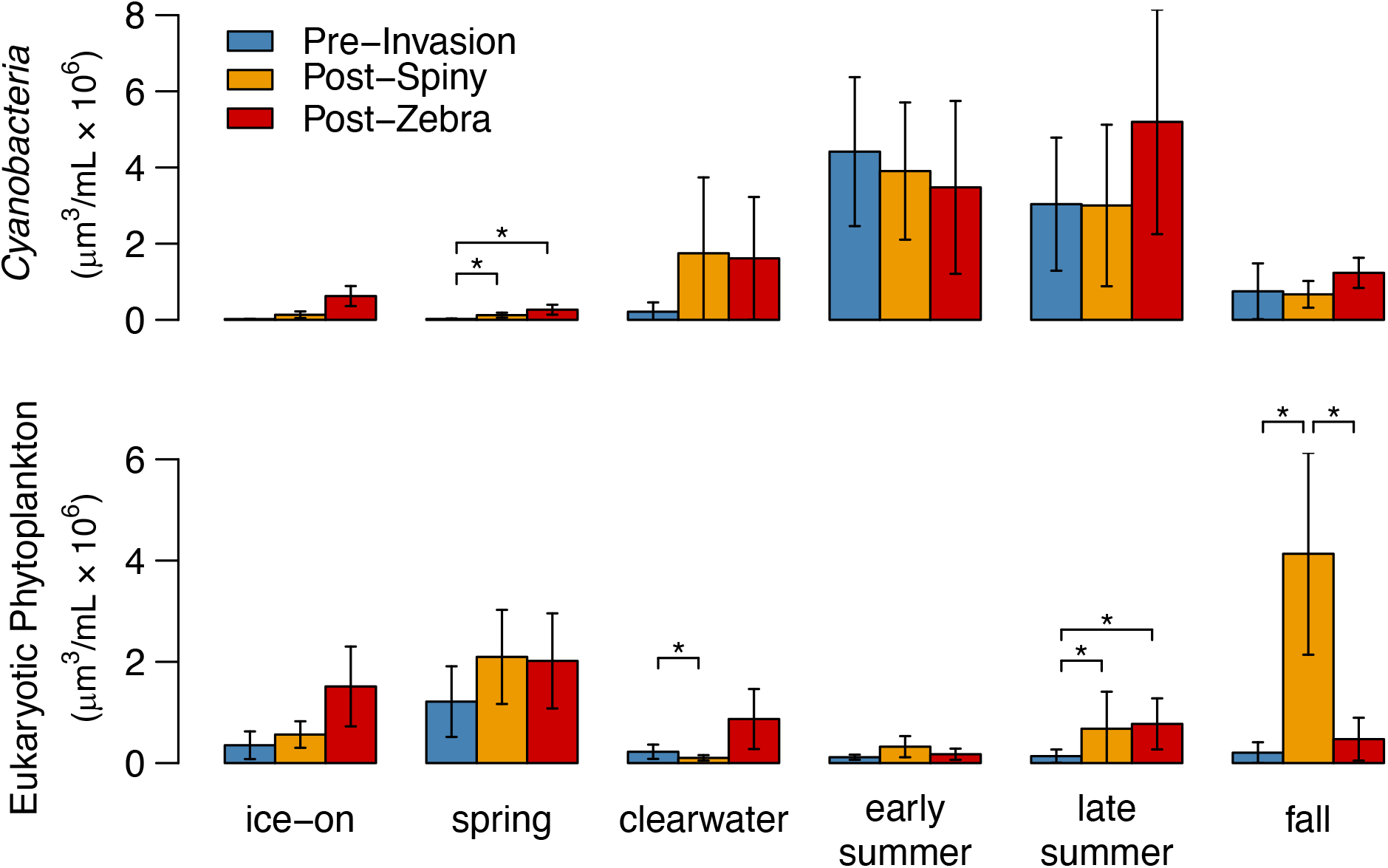
Phytoplankton biovolume. The biovolume of *Cyanobacteria* and eukaryotic phytoplankton in each season, broken down by invasion status.

**Figure S2.**
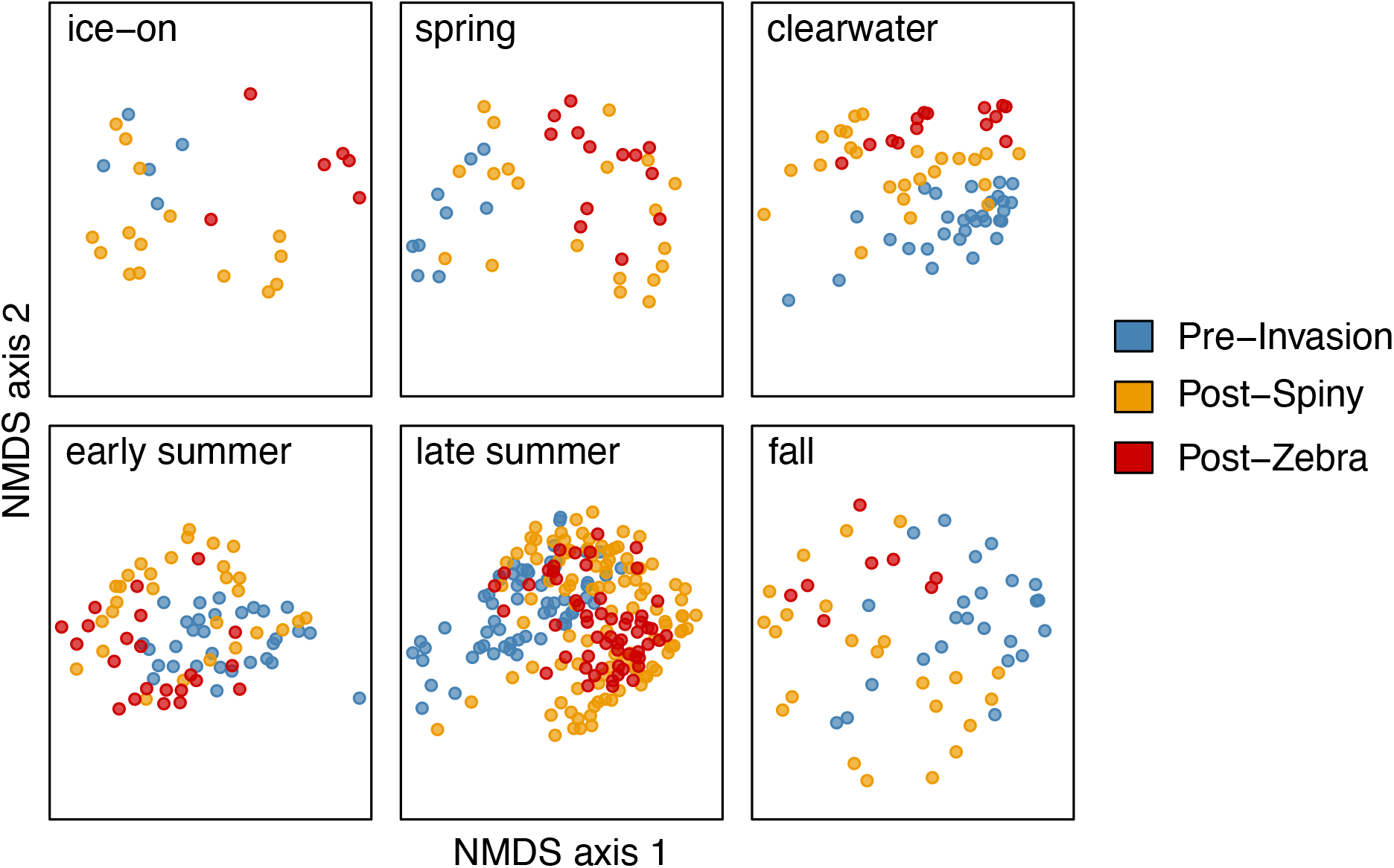
NMDS plots of invasion impacts by season. Panels correspond to different seasons, points represent sample dates included in each season, and colors indicate the sample date’s invasion status. NMDS analysis was performed on CLR-transformed data.

**Table S1.**
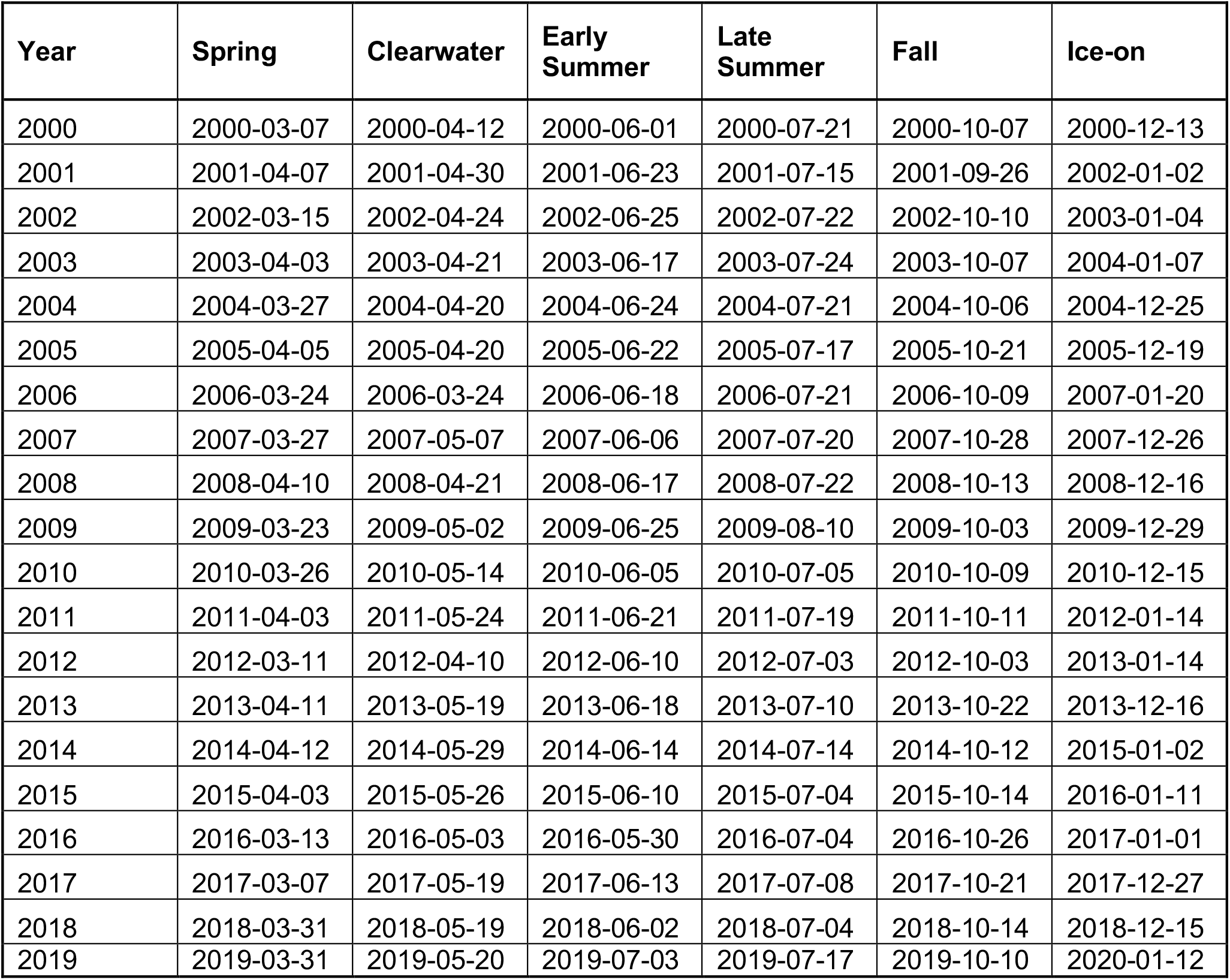
Season start dates.

**Table S2.** Cyanotoxin concentrations. *Larger than a single page*.

**Table S3.**
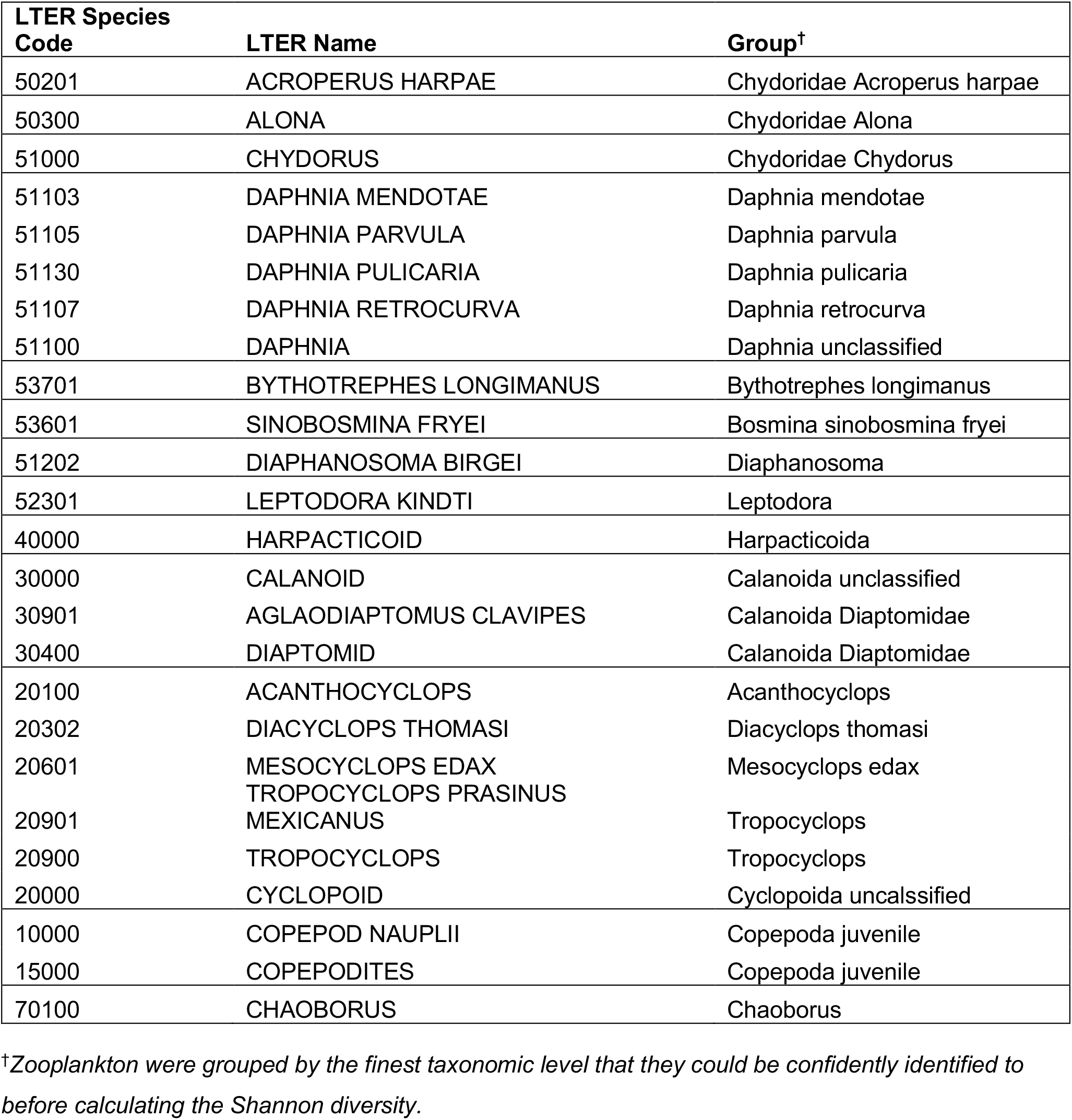
Zooplankton grouping for diversity calculation.

**Table S4.** Differentially abundant OTUs. *This table is larger than a single page*.

**Table S5.** AIC table for models predicting *Cyanobacteria* phenology.

| Hot<sup>1</sup> | Ice<sup>2</sup> | Pload<sup>3</sup> | SWF<sup>4</sup> | ZM<sup>5</sup> | AICc | delta | weight | df |
| --- | --- | --- | --- | --- | --- | --- | --- | --- |
| 0.31 |  |  | -36.52 | -37.19 | 163.22 | 0.00 | 0.30 | 5 |
|  |  |  | -34.27 | -35.83 | 163.29 | 0.07 | 0.29 | 4 |
| 0.51 |  | 0.53 | -35.83 | -42.24 | 163.77 | 0.55 | 0.23 | 6 |
|  |  | 0.09 | -33.91 | -36.57 | 166.92 | 3.70 | 0.05 | 5 |
|  | -0.01 |  | -34.15 | -36.05 | 167.03 | 3.82 | 0.05 | 5 |
| 0.33 | 0.06 |  | -37.17 | -36.26 | 167.39 | 4.18 | 0.04 | 6 |
| 0.50 | -0.02 | 0.55 | -35.52 | -42.83 | 168.90 | 5.69 | 0.02 | 7 |
|  |  |  | -46.21 |  | 170.74 | 7.52 | 0.01 | 3 |
|  | -0.04 | 0.12 | -33.43 | -37.52 | 171.23 | 8.01 | 0.01 | 6 |
| 0.26 |  |  | -48.47 |  | 172.55 | 9.33 | 0.00 | 4 |
|  | 0.11 |  | -46.56 |  | 173.60 | 10.38 | 0.00 | 4 |
|  |  | -0.12 | -46.36 |  | 173.87 | 10.65 | 0.00 | 4 |
| 0.33 | 0.18 |  | -49.58 |  | 175.23 | 12.01 | 0.00 | 5 |
|  |  |  |  | -57.25 | 175.84 | 12.62 | 0.00 | 3 |
| 0.31 |  | 0.13 | -48.70 |  | 176.20 | 12.98 | 0.00 | 5 |
|  | 0.14 | -0.22 | -46.95 |  | 176.98 | 13.76 | 0.00 | 5 |
|  |  | 0.28 |  | -58.78 | 178.59 | 15.37 | 0.00 | 4 |
| 0.16 |  |  |  | -58.68 | 178.68 | 15.46 | 0.00 | 4 |
|  | -0.13 |  |  | -58.76 | 178.69 | 15.48 | 0.00 | 4 |
| 0.34 | 0.17 | 0.03 | -49.62 |  | 179.60 | 16.39 | 0.00 | 6 |
| 0.39 |  | 0.62 |  | -64.14 | 180.60 | 17.38 | 0.00 | 5 |
|  | -0.21 | 0.43 |  | -62.18 | 181.34 | 18.13 | 0.00 | 5 |
| 0.13 | -0.10 |  |  | -59.65 | 182.18 | 18.96 | 0.00 | 5 |
| 0.39 | -0.21 | 0.78 |  | -67.46 | 183.91 | 20.69 | 0.00 | 6 |
|  |  |  |  |  | 186.74 | 23.52 | 0.00 | 2 |
|  | 0.04 |  |  |  | 189.57 | 26.35 | 0.00 | 3 |
| -0.03 |  |  |  |  | 189.58 | 26.37 | 0.00 | 3 |
|  |  | -0.02 |  |  | 189.59 | 26.37 | 0.00 | 3 |
|  | 0.05 | -0.05 |  |  | 192.82 | 29.60 | 0.00 | 4 |
| -0.02 | 0.04 |  |  |  | 192.82 | 29.61 | 0.00 | 4 |
| -0.06 |  | -0.07 |  |  | 192.83 | 29.61 | 0.00 | 4 |
| -0.05 | 0.05 | -0.09 |  |  | 196.56 | 33.35 | 0.00 | 5 |

<sup>1</sup>Days with average epilimnion temperature $\geq 23^{\circ}\text{C}$ .
<sup>2</sup>Previous winter ice duration in days.
<sup>3</sup>Mean annual daily phosphorus loading.
<sup>4</sup>Spiny water flea invasion status.
<sup>5</sup>Zebra mussel invasion status.

**Table S6.**
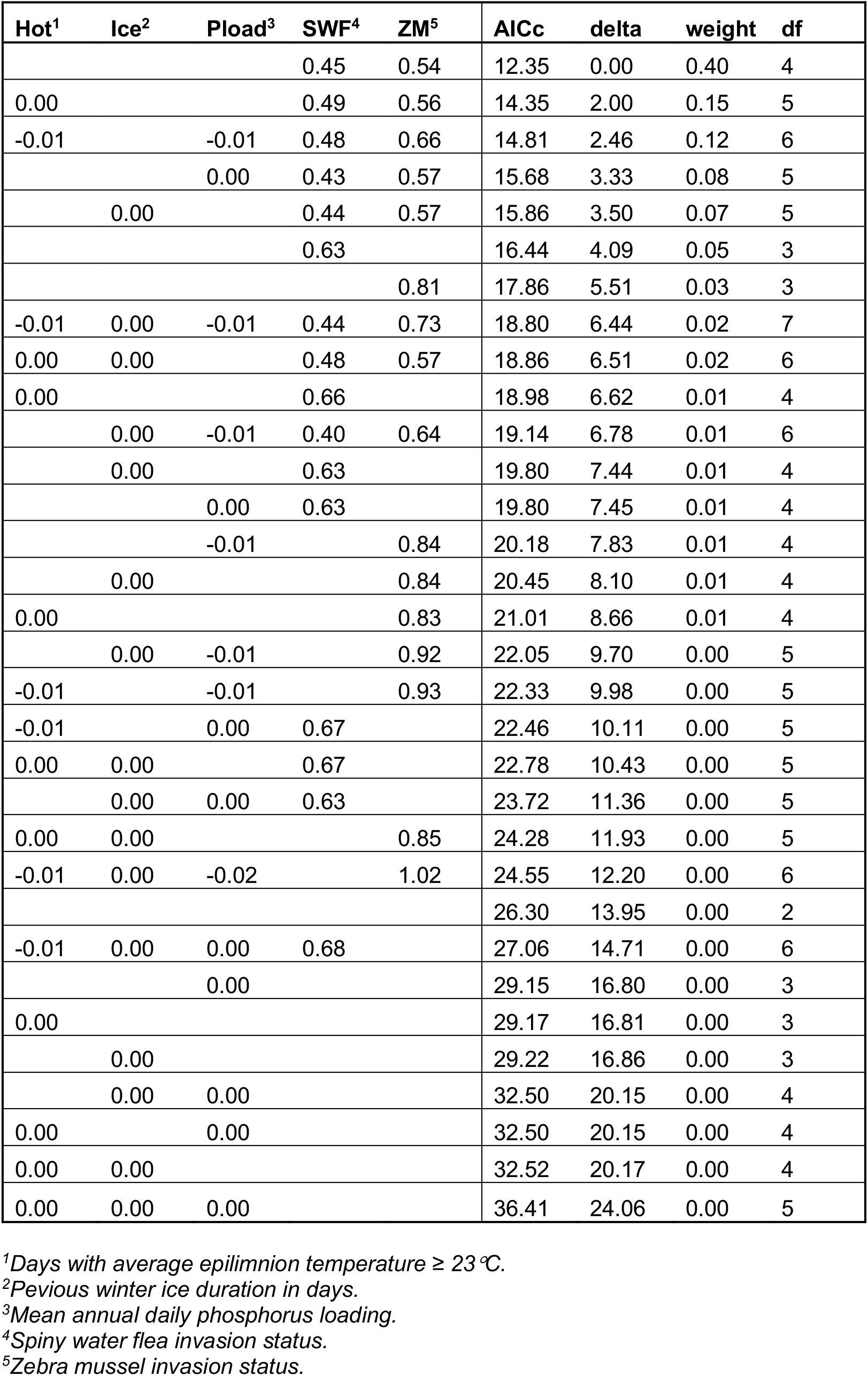
AIC table for models predicting *Frankiales* phenology.

